# Modulatory feedback determines attentional object segmentation in a model of the ventral stream

**DOI:** 10.1101/2023.01.19.524712

**Authors:** Paolo Papale, Jonathan R. Williford, Stijn Balk, Pieter R. Roelfsema

**Affiliations:** Department of Vision & Cognition, Netherlands Institute for Neuroscience (KNAW), 1105 BA Amsterdam, Netherlands; STR, Woburn, MA, USA; Department of Integrative Neurophysiology, VU University, De Boelelaan 1085, 1081 HV Amsterdam, Netherlands; Department of Neurosurgery, Academic Medical Centre, Postbus 22660, 1100 DD Amsterdam, Netherlands; Laboratory of Visual Brain Therapy, Sorbonne Université, INSERM, CNRS, Institut de la Vision, 17 rue Moreau, Paris, F-75012, France

## Abstract

Studies in neuroscience inspired progress in the design of artificial neural networks (ANNs), and, vice versa, ANNs provide new insights into the functioning of brain circuits. So far, the focus has been on how ANNs can help to explain the tuning of neurons at various stages of the visual cortical hierarchy. However, the role of modulatory feedback connections, which play a role in attention and perceptual organization, has not been resolved yet. The present study presents a biologically plausible neural network that performs scene segmentation and can shift attention using modulatory feedback connections from higher to lower brain areas. The model replicates several neurophysiological signatures of recurrent processing. Specifically, figural regions elicit more activity in model units than background regions. The modulation of activity by figure and ground occurs at a delay after the first feedforward response, because it depends on a loop through the higher model areas. Importantly, the figural response enhancement is enhanced by object-based attention, which stays focused on the figural regions and does not spill over to the adjacent background, just as is observed in the visual cortex. Our results indicate how progress in artificial intelligence can be used to garner insight into the recurrent cortical processing for scene segmentation and object-based attention.

**Author Summary:** Recent feedforward networks in artificial intelligence provide unmatched models of tuning of neurons in the visual cortex. However, these feedforward models do not explain the influences of object-based attention and image segmentation on neuronal responses, which rely on feedback interactions between cortical regions that are not included in the feedforward networks. In particular, the role of feedback connections from higher brain regions that modulate neural activity in lower cortical regions has not yet been studied extensively so that we still lack an *in silico* model of the role of these connections. Here, we present a biologically plausible neural network that successfully performs image segmentation and can shift object-based attention using modulatory feedback connections. The model evolved representations that mirror the properties of neurons in the visual cortex, including orientation tuning, shape-selectivity, surround suppression and a sensitivity to figure-ground organization, while trained only on a segmentation task. The new model provides insight into how the perception of coherent objects can emerge from the interaction between lower and higher visual cortical areas.

## Introduction

Objects are the building blocks of our perception. We perceive a visual scene as a layout of objects in the physical space. We tend to focus on one object at a time when scanning a visual scene, either overtly while moving our eyes or covertly by shifting attention (O’Craven et al., 1999). This process, called object-based attention, is essential to group all the visual features that surround us into the objects with which we interact (Behrmann et al., 1998; Duncan, 1984; Kramer et al., 1997; Roelfsema, 2023). When we select a particular object for action, object-based attention is the mechanism that helps identifying the portions of the scene that belong to that object and segregating it from other objects and the background. If subjects grasp and manipulate small objects, attentional selection can occur at the required precise spatial scale (Roelfsema, 2006; Roelfsema and Houtkamp, 2011).

Although it may appear effortless to us, object identification and object-based attention in the primate visual system rely on elaborate computations in a hierarchy of cortical regions. The input of the retina first reaches the primary visual cortex (V1) where neurons have small receptive fields (RFs) and are tuned to simple features. Information is then propagated through feedforward connections to higher-order regions in the occipital cortex (areas V2 and V4) and onward to inferotemporal (IT) and frontal (FC) cortices (van Essen et al., 1992). In the higher areas the neurons become selective to more complex features, such as object parts and object identity and they have larger receptive fields.

However, if the task is to segment the scene, the computations carried by feedforward connections are only half of the story (Kar et al., 2019; Lindsay and Miller, 2018; Thorat et al., 2019). After reaching the higher visual areas, visual information is also carried back to the lower areas through feedback connections (Buffalo et al., 2010; Felleman and van Essen, 1991) (Fig 1A). Whereas feedforward connections drive the initial response of neurons and shape neural selectivity, feedback connections modulate the level of neural activity. Feedback connections enhance the neuronal activity elicited by relevant figural regions over the activity elicited by the background, a process called figure-ground modulation (Lamme, 1995; Poort et al., 2016; Roelfsema and de Lange, 2016; Zipser et al., 1996). As a result, all the features of objects are enhanced in lower visual brain areas, such as V1, at a high spatial resolution. Furthermore, feedback connections mediate the influence of top-down attention that amplify neuronal activity elicited by task-relevant objects (Roelfsema, 2023, 2006; Spitzer et al., 1988; Vecera and Farah, 1997).

**Fig 1.**
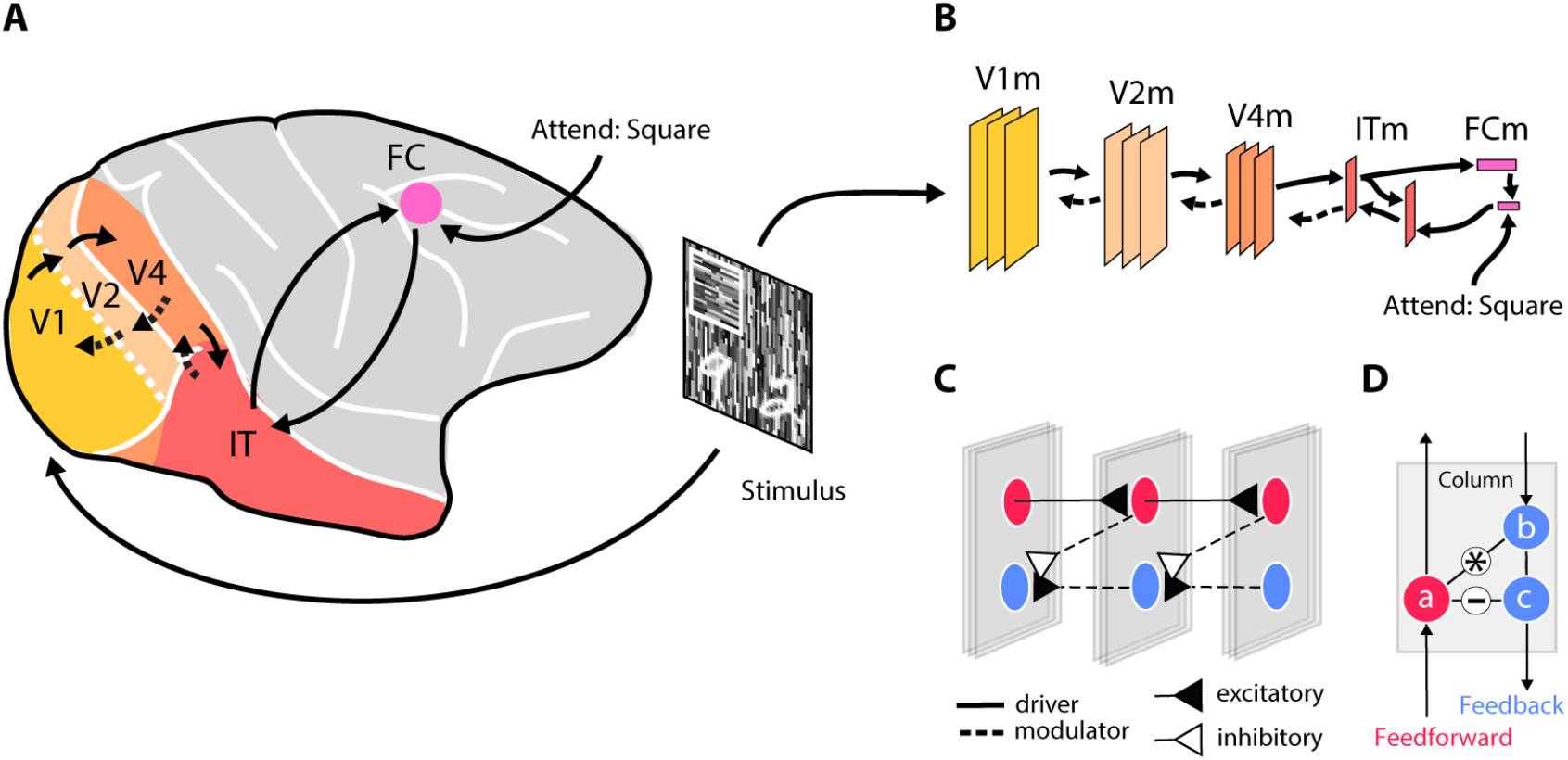
A neural network to model the primate visual cortex. A) Hierarchically organized visual cortical areas that perform increasingly complex transformations of the visual input. V1 registers the information coming from the retina. Downstream regions (V2, V4) process information further and object-selective neurons in IT categorize the objects in a scene and send this information to the frontal cortex (FC) that represents behavioral goals. The frontal cortex sends attentional signals through feedback connections that modulate the activity of earlier brain regions, resulting in the segmentation of task-relevant objects at a high spatial resolution. B) In the model, the stimulus enters in area V1 (V1m) and feedforward connections (solid arrows) activate downstream regions that classify the objects. The network can select one of the object classes and use feedback connections for class-specific object segmentation in V1m. C) The pattern of connections between V1m, V2m and V4m. The network contains units that propagate information in the feedforward direction (red) and other units (blue) that propagate information in the feedback direction. The signal flowing in the feedback direction is based on a comparison (subtraction) of the activity of feedback and feedforward units. D) Schematic of the computations within one of the layers. *a* is the activity of feedforward unit, which propagates activity to the next higher layer, gates information flow of unit *b* (a multiplicative influence; *) and inhibits *c* (-) of the feedback pathway.

At first sight, many of these neurophysiological observations may seem disconnected, because previous studies focused on only one or a few of these feedback influences (Carandini et al., 2005), or used models with connectivity that was largely handcrafted (Craft et al., 2007; Deco and Rolls, 2004; Hamker, 2005; Van Der Velde and De Kamps, 2001). We therefore sought to integrate previous findings into a coherent framework, capitalizing on the recent advances in the development of artificial neural networks (ANNs) (Echeveste et al., 2020; Fox et al., 2023; Serre, 2019).

State of the art ANNs can achieve human level performance on many complex visual tasks, ranging from object classification to segmentation (Kirillov et al., 2023; Serre, 2019). For example, breakthroughs were achieved in the problem of semantic segmentation, where the goal is to determine the pixels that belong objects of specific categories (Kirillov et al., 2023; Li et al., 2018). ANNs for semantic segmentation have features in common with segmentation processes in the brain, by relying on two pathways. The first pathway starts with units representing pixels and end with units coding for object categories through a hierarchy of layers with increasing RFs and more complex tuning. The second pathway propagates activity in the opposite direction, from object category back to pixels, so that tuning becomes simpler and RF size decreases during successive processing steps (Hong et al., 2015). The selectivity of units in the feedforward pathway of ANNs, resembles the selectivity of neurons in the visual brain (Cadena et al., 2019; Guclu and van Gerven, 2015; Yamins et al., 2014). However, the resemblance of the ANN feedback pathways to neurobiology is more limited. Firstly, in most ANNs units of the feedback pathway are independent from those of the feedforward pathway (e.g., Hong et al., 2015), whereas many neurons in the brain are influenced by both feedforward and feedback connections (e.g., Lei et al., 2021). Secondly, in the visual cortex, neurons that are influenced by feedback and those that are mainly feedforward tend to segregate into different layers of the cortical column. The feedback influences are weak in cortical layer 4, which is the input layer of cortex, and stronger in the superficial and deep layers (Keller et al., 2020; Kirchberger et al., 2023; Self et al., 2013; van Kerkoerle et al., 2017). Interestingly, neurons with different degrees of sensitivity to feedback effects reside in the same column and have a similar tuning. Thirdly, the feedback pathway of ANNs drives the units, whereas feedback connections in the brain are mostly modulatory. This means that feedback connections usually do not directly activate the neurons but amplify or decrease the feedforward response. Some of these feedback influences are multiplicative, with a stronger influence on neurons that are well driven by feedforward input than on neurons that are not (Poort et al., 2016, 2012; Roelfsema et al., 2002; Self et al., 2012; Treue and Martínez Trujillo, 1999).

Hence, the brain processes underlying image segmentation and, in particular, semantic segmentation have remained unclear. How do the feedback pathways of the brain, which are modulatory and intermingled with the feedforward pathways, achieve object segmentation? To answer these questions, we built a biologically inspired ANN to examine figure-ground segregation, semantic segmentation and the role of selective attention (Poort et al., 2012). In our model, feedforward and feedback influences occur within the same cortical column and feedback is modulatory (Keller et al., 2020; Self et al., 2013; van Kerkoerle et al., 2017). The network comprised a hierarchy of areas modeled after the hypothetical brain regions involved in object processing – denoted here with an ‘m’ suffix – from V1m to FCm (Fig 1B). We included a feedforward pathway, driving the activity in successively higher areas for object identification, and a feedback pathway that modulated neuronal activity (Fig 1C,D) to highlight the figural image regions if they were attended.

The model produced an accurate segmentation of attended objects by increasing the firing rate elicited by image elements in V1m, representing the relevant figure at a high spatial resolution, whereas the activity elicited by nearby background elements was suppressed, just as is observed in the visual cortex of monkeys (Poort et al., 2012; Self et al., 2019). When we examined the tuning of model units we observed similarities to the neurophysiology, indicating that the model enables new insights in how feedforward and feedback connections jointly determine the activity of neurons in the visual cortex.

## Materials and Methods

### Model Architecture

Fig 1B provides an overview of the model, which consists of several model areas, which loosely reproduce the ventral stream of the visual cortex of the primate. Our architecture was inspired by an ANN (Hong et al., 2015) that performs semantic segmentation in separate feedforward and feedback pathways, i.e. its goal is to assign an object class to pixels of the input image (Fig. 2A). The architecture introduced by Hong et al. (2015) achieved this by first mapping pixels into object classes in a feedforward network. It then selected one of the classes at a time for a given image and retrieved the pixels belonging to that class in an additional feedforward network that mapped classes back onto pixels. Hence, this ANN carried out an object-based attention task in a purely feedforward network. Our model, instead, used feedforward connections that are driving and feedback connections that are modulatory. It can therefore provide insight into the different phases of activity of neurons in the visual cortex.

**Fig 2.**
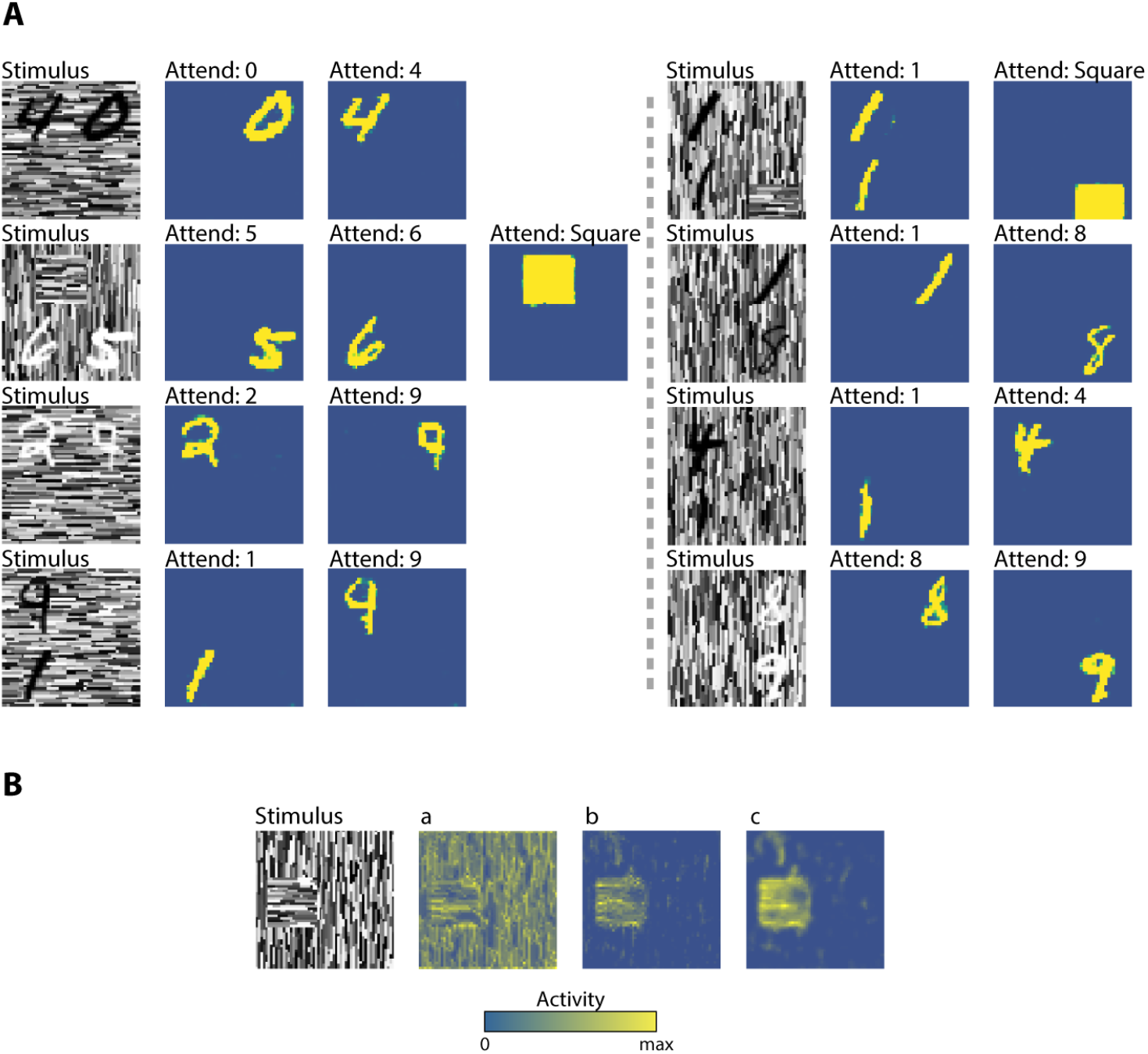
Modulatory feedback achieves accurate segmentations of attended objects in cluttered displays. A) Eight example stimuli and the segmentations achieved by the model. Each row shows an example stimulus and the segmentation generated by the model when instructed to attend to a specific object class (blue, low activity level denoting background; yellow, high activity level labeling the segmented object). The segmentations in V1m are accurate, demonstrating that modulatory feedback connections enable a selective segmentation of the attended object category. B) The internal representation of an example stimulus (left), show how the activity of V1m feedforward units “*a”* (middle left), is subtracted from the modulatory feedback “*b”* (middle right) coming from a downstream region, to generate the final modulated response “*c”* (right).

The model comprised two main processing streams, a feedforward pathway that drives the neurons and a feedback pathway that is modulatory. The feedforward pathway took an image as input and produced a classification vector that indicated the object classes that were detected in the image as output. This feedforward pathway was not modulated by feedback, akin to what observed in the input layer 4 of area V1 (Keller et al., 2020; Self et al., 2013; van Kerkoerle et al., 2017). The feedback pathway contained units that were influenced by the feedforward units in the same area and that were also modulated by feedback connections, as will be detailed below. The feedback pathway started with the classification vector at the highest levels of the feedforward pathway, which it combined with a one-hot attention vector that specified a target category (e.g. “attend the texture square” in Figure 1A). The goal of the feedback pathway was to highlight all the low-level features elicited by the target object with enhanced neuronal activity.

### Feedforward pathway

The network layers were grouped into areas that roughly correspond to visual cortical areas. All the layers within a given area had the same size. Areas V1m, V2m and V4m had 20 features and 56×56 pixels, 50 features and 28×28 pixels, and 100 features and 14×14 pixels, respectively, resulting in larger RFs in higher areas (i.e. 2×2 pixels in V1m, 4×4 pixels in V2m and 8×8 pixels in V4m). Each area had three convolutional layers with skip connections bypassing the convolution followed by batch normalization and a ReLU function. The output of each area was downsampled by a strided convolution (stride of 2 and a 2×2 kernel size). The down-sampled output from V4m (100 features by 7×7 pixels) was passed to a fully connected layer (ITm: 500×1×1), which was then passed to another fully connected layer (FCm: 12×1×1), which had one unit for each of the 12 object classes in our task. This layer used a sigmoid activation function (i.e. softmax) to estimate the likelihood that the various classes appeared in the stimulus. Multiple object classes could appear at the same time.

### Feedback Pathway

The interactions between the feedforward and feedback pathway ensured that the feedback from the selected shape in FCm modulated the appropriate lower-level units. A 12×1×1 layer encoded the object that should be attended (the “attend” label in Fig. 1B). This attention signal was multiplied (elementwise) with the classification vector from the feedforward pathway. A series of fully connected layers were used in the feedback pathway, mirroring the feedforward pathway, followed by a “deconvolution” (transposed convolution) layer that produced output with dimensions of 100×7×7. This output was then concatenated with the corresponding 100×7×7 layer of the feedforward to produce a 200×7×7 tensor, which was passed to a 1×1 convolution reducing the tensor back to 100×7×7.

Each unit in the feedforward pathway had a corresponding (deconvolutional) unit in the feedback pathway in V1m, V2m and V4m, which can be conceived of as being part of the same cortical column (Fig. 1D). The feedback connections did not drive units in these areas, but they modulated the activity, in accordance with the effects of attention in the brain (e.g., Self et al., 2012). In our model the modulatory feedback interaction was implemented as:

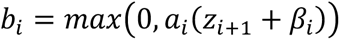

where *a*_*i*_ is the non-negative firing rate of a feedforward neuron in layer *i, z*_*i*+1_ is the feedback received from a higher layer via a strided convolution, *β*_*i*_ is a learned parameter, and *b*_*i*_ is the activity of a unit of the feedback pathway (Fig. 1D, Fig. 2B). The feedback signal that is propagated to the next lower layer, is the difference between the activity of the feedforward and feedback units and it constitutes a relatively pure segmentation signal:

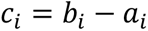

### Stimulus sets

Based on the MNIST dataset (Yann LeCun et al., 2010), we created an augmented stimulus set, *SegMNIST* (Fig. 2A). We followed recent approaches to make MNIST-like stimulus sets more challenging (Ernst et al., 2019; Michaelis et al., 2018; Spoerer et al., 2017; Thorat et al., 2021) and casted it as an object localization task, where the relevant shape could appear at various locations in the image. The size of the image was increased to twice the size of the MNIST image (56×56 pixels instead of 28×28) and MNIST digits were translated, scaled, and positioned on top of a texture. To gain insight in figure-ground segregation processes, we also added textured squares and rectangles at different rotation angles with an orientation that differed from the background, which have been used in previous monkey experiments on texture segregation (Poort et al., 2012; Self et al., 2012)

### Training of the feedforward and feedback pathways

The feedforward pathway was trained to classify objects in the textured and expanded SegMNIST stimuli using Adam (Kingma and Ba, 2015) with a learning rate of 0.001, L2-regularization (weight decay = 0.0005), and as loss a sigmoid combined with binary cross-entropy (pytorch’s *BCEWithLogitsLoss*) in 2,500 epochs (batch size = 1024). There were 12 object classes; a square, a rectangle, and the 10 digits, and there were one to three objects in each image. The task of the network was to determine which object classes were present in the image. We trained the feedback pathway once the weights of the feedforward pathway were fixed. The feedback pathway was trained on object segmentation using Adam (Kingma and Ba, 2015) with learning rate of 0.001, L2-regularizations (weight decay = 0.0005), and as loss a sigmoid combined with binary cross-entropy (pytorch’s *BCEWithLogitsLoss*) in 2,500 epochs (batch size = 256). The objective was to maximize the number of pixels in the image that were correctly classified (with activity 1 for pixels belonging to the attended class and 0 elsewhere). Because of the imbalance in the number of pixels in objects and the background, we ignored background pixels when computing the error, because it would have biased the error and led to poor generalization. The network was trained using one GPU.

### Simulating the activity propagation in V1m

To evaluate the influence of attentional selection on the activity across time, we allowed the feedforward activity to propagate to layers in successive time steps. The initial feedback modulation could proceed in lower layers while the feedforward activity was propagating to the higher layers of the network (Fig. 3). We simulated a single feedforward pass and a single feedback pass but did not model full recurrence. Our simulations address the situation in which the subject aims to select (attend) a specific shape. We initiated the network at the activity level elicited by a white noise stimulus and activated the attentional template of the to-be-selected shape in FCm. We then presented the stimulus and the activity propagated to the next higher and lower layer on every time step. We presented 1,000 stimuli (with the model attending to either figures or digits) and averaged the responses. The time-courses shown in Figure 3B,D were up-sampled (to 160 steps) and smoothed with a 3^rd^ order Savitzky–Golay filter (7 timestep window) for visualization purposes.

**Fig 3.**
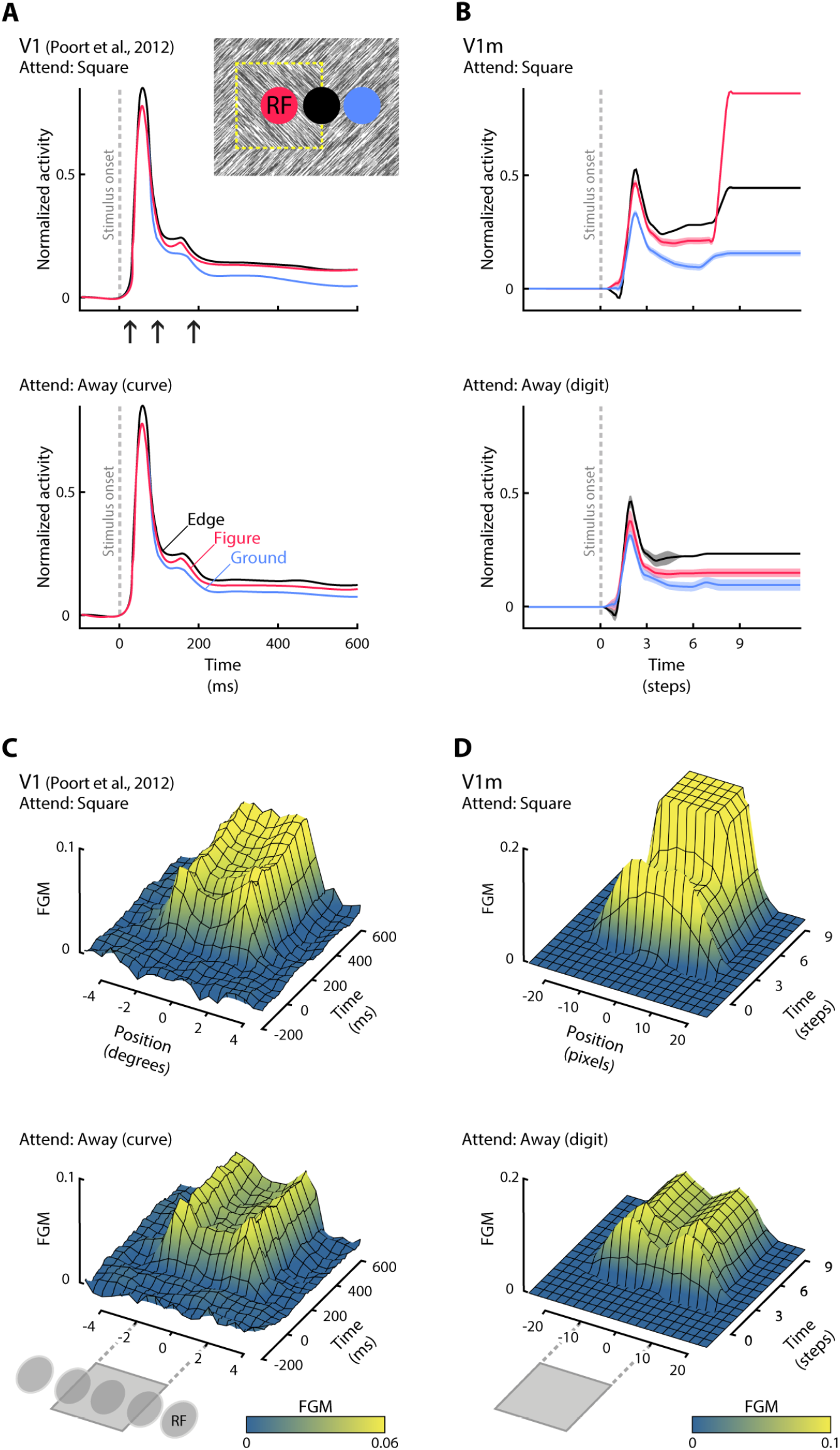
V1m evolves figure-ground modulation by selective attention as observed in monkey V1. A) Activity of V1 neurons in the study of Poort et al. (2012), in which monkeys were cued to either detect texture-based square figures (top) or to perform another task in which they did not attend the figures. V1 activity depended both on whether the texture elements falling in RF were part of the figure, the ground or the edge between figure and ground and also on attention. There were three different phases of activity: a first transient response (top, left arrow), a later phase in which figures elicited more activity than grounds (middle arrow) and an even later phase in which attention increased the figure response (right arrow; compare the distance between red and blue curves in the upper and lower panels). B) Response of feedback units in V1m when attention is directed to the figure (top) or to one of the digits (bottom). The activity was propagated for one connection at a time. Dashed regions represent SEM (across 1,000 stimuli with different textures). C) Space-time profile of figure-ground modulation (FGM: difference between the figure and background response) for V1 neurons when the monkey attended to the figure (top) and or attended away (bottom). The horizontal axis represents the position of the RF, which either fell on the figure, the edge or the background. D) Space-time profile of V1m units when the model attended the square (top) or one of the digits (bottom).

### Code availability

Model and analyses were implemented using custom Python code and NumPy (van der Walt et al., 2011), SciPy (SciPy.org) and PyTorch (Paszke et al., 2019). The code to generate the stimuli is available at the SegMNIST Github repository: http://github.com/williford/seg-mnist-dataset. The code to train the model and reproduce the results is available at the Github repository of the project [available upon paper acceptance]. A Google Colab of the project with the model results and all units is available at https://colab.research.google.com/drive/13O0-4uYq3l1tZupVn-NLP8C-WvakajCG?usp=sharing.

## Results

### Modulatory feedback connections achieve accurate segmentations of attended objects

We trained an ANN to recognize and segment the digits 0 through 9, textured squares and rectangles in a display with a background of texture elements with an orientation that differed from the foreground squares and rectangles. First, we trained the feedforward path of the network to recognize these shapes. Then we trained the feedback pathway to attend and segment one of the shapes from the background. In this training phase, we used a task that was inspired by Poort et al. (2012), who trained monkeys to either detect texture-based squares similar those of Fig. 2 or to perform another task in which they could ignore these texture-defined figures.

A few example stimuli and the model’s segmentation results are shown in Fig. 2A. The stimuli (left column) were particularly challenging because the edges of the digits blended with the lines of the texture in the background and the squares and rectangles were defined by a difference in the orientation of the texture elements. Despite these challenges, the classification accuracy of feedforward path of the model was nearly perfect. Specifically, the activation of the FCm units reflected the objects classes with a mean cross-validated absolute error of 0.023 (this measure ranges between 0 when all objects are correctly classified in each image and 12 if none of the objects are correctly classified).

We next examined if and how neurons of the feedback pathway were influenced by the presence of figure-ground stimuli and the shape that had been selected in FCm. The feedback pathway caused highly accurate segmentations. Responses of feedback units to images regions of the attended category were enhanced over the responses of units that responded to the background and over the responses elicited by non-attended objects (yellow regions in Fig. 2A denote attended regions, which have been labeled with enhanced activity; blue regions are labeled as background). Consider, for example, the left stimulus in the second row of Fig. 2A that contained a square and two digits, a ‘5’ and a ‘6’. When the square was attended, by switching on the corresponding unit in FCm, the activity of V1m units with receptive fields on the square was enhanced. This response enhancement was confined to the square at a high spatial precision and it did not spill over into the background. However, if attention was directed to one of the digits, the responses to the square were weaker and the response enhancement was confined to the pixels of the appropriate digit. These highly accurate segmentations occurred for most of the images, so that the mean cross-validated pixel-wise absolute error was 0.0015 (the error ranges between 0 when all object pixels correctly segmented and 1 if none of the object pixels is correctly segmented in each image; this measure does not consider background pixels). Furthermore, the proportion of background elements that were erroneously labeled as the attended class was very small (<0.1%).

These results demonstrate that modulatory feedback connections can explain the selective labeling of low-level image elements of an attended object category with enhanced neuronal activity, just as is observed in the primary visual cortex of monkeys (Lamme, 1995; Poort et al., 2012; Self et al., 2019; Zipser et al., 1996). When we examined the tuning of model units to shape and orientation, we also observed strong similarities to the properties of neurons in the visual cortex of monkeys, and the same held true for surround suppression (Fig. S1,S2). Interestingly, tuning to the shape of objects was stronger in the feedback pathway in V4m and V2m than in the feedforward pathway, indicating that feedback connections carried information about the identity of the attended objects from higher regions (ITm and FCm) to lower regions (Fig. S2).

We next compared the time-course of the activity of model units to the neurophysiological results of Poort et al. (2012). In that study, monkeys either attended texture-defined square figure, similar to the stimuli shown in Fig. 2A, or performed another task in which they ignored the square. The main neurophysiological results are summarized in Fig. 3A, showing the time-course of the response of V1 neurons with a RF on the center of the square figure (red trace), the edge between the square and the textured background (black trace) or the background (blue trace). The activity started around 40ms after the presentation of the stimulus, when feedforward activations drive the V1 neurons. During this early response phase, the activity depends on the information in the RF, and after a short delay the activity of neurons with a RF on the edge between figure and ground is enhanced over the activity of neurons with an RF on the background. During a later response phase, around 100ms after the stimulus onset, the activity elicited by image elements that belong to the center of the figure start to be also enhanced over the activity elicited by background elements. Finally, approximately 200ms after onset, the influence of attention to the figure (Fig. 3A, top) kicks in. Attention to the figure causes a further enhancement of the activity of V1 neurons with a RF on figure region, compared to when monkeys ignore the figure (Fig. 3A, note that the separation between black and red traces is more pronounced in the lower panel).

We examined the activity of V1m units in a version of the model in which we propagated activity in the feedforward and feedback directions by one connection at a time. The activity profile of V1m units was similar to the activity of neurons in monkey V1. The model units exhibited a first, transient peak response that was elicited by the appearance of texture elements in the RF (Fig. 3B). The early response elicited by the edge (black in Fig. 3B), was higher than that elicited by the figure and ground and the response elicited by the figure was stronger than that elicited by the background. At a later phase, attention to the textured square enhanced the activity evoked by the figure further (Fig. 3B, top), compared to when attention was directed to one of the digits (Fig. 3B, bottom). The influence of attention on neuronal activity in the model feedback pathway was stronger than in monkey V1 and activity elicited by the center of the figure exceeded the activity elicited by the edge. This difference between the model and the neurophysiological results is explained by the model’s training objective, which enforced binary segmentation where figural regions reach a target activity of 1. However, it is of interest that the order of the effects of figure-ground organization and attention was similar to that in monkey V1.

Figure 3C,D compares the spatial profile of the figure-ground modulation (FGM), which is difference in activity elicited by the figure and the background (subtraction of the blue curve from the red/black curve in Fig. 3A,B), in monkey V1 (Fig. 3C) and in the feedback pathway in model area V1m (Fig. 3D). The monkeys saw a figure with a size of 4 degrees and the size of the V1 RFs was less than 1 degree. In the experiment with monkeys the location of the figure was varied so that the neurons’ RFs could either fall on the figure, the background or the edge. It can be seen that FGM was confined the figure representation. If the monkey did not attend the figure, FGM was stronger at the boundary between figure and background than in the center of the figure. If the monkey attended the figure, however, FGM in the center of the figure was more pronounced and reached the same level as that at the boundary.

Many of these observations also held true for the feedback units in V1m of the model, where FGM was also confined to the figure. If attention was directed to one of the digits, FGM was strongest at the boundaries of the square and weaker in the center of the figure. However, if the model attended the textured square, activity was strong in the center of the figure. In the model, the activity reached a higher plateau, which is related to the training procedure in which the target activity level in V1m was 1, as was described above.

## Discussion

In this work, we built a biologically inspired ANN, implementing a model of the primate object processing pathway, which was composed of a driving feedforward pathway and a modulatory feedback pathway. We demonstrated that this simple architecture could shift selective attention to all the low-level features that belong to the same object, even it is embedded in a cluttered texture stimulus. We compared the activity of model units to the responses of neurons in the visual cortex of monkeys. Firstly, the model reproduced the spatial profile of neuronal activity in the primary visual cortex during image segmentation tasks, with a strong enhancement of the representation of image elements of figures relative to the background, even though the image elements of figure and ground were, on average, the same. Secondly, the time-course of activity of model units in V1m resembled the time-course of the response of V1 neurons. Activity started with a transient response, followed by figure-ground modulation and the influence of attention was expressed at an even later time-point, because feedforward processing had to propagate to FCm, which then had to reach back to V1m. The delayed attentional influence is in accordance with neurophysiological evidence about comparable top-down influences (e.g., Bichot et al., 2019, 2015; Moore and Fallah, 2004; van Vugt et al., 2018). However, the attentional influence in V1m was stronger than in monkey V1. A likely reason for this difference is that our model was trained on a binary segmentation task with a high target value of attended pixels, whereas V1 of animals contributes to many more tasks (Roelfsema and de Lange, 2016), which is a limitation of our model. Thirdly, we found qualitative similarities between the properties of neurons in the visual cortex and in the model, including orientation tuning, surround suppression and shape-selectivity (Figs. S1,S2). It is of interest that tuning to object identity was strongest in the feedback pathway in V4m and V2m, indicating that it was based on feedback from the higher model areas (Fig. S2). These results thereby provide insight why object classes can be decoded from the activity of early visual neurons in the primate brain (e.g., Muckli et al., 2015; Papale et al., 2023, 2021, 2020, 2019; Williams et al., 2008).

Previous modeling studies addressing the neuronal mechanisms of image parsing and texture segregation in the visual cortex used hand-crafted connectivity schemes and activation functions (Poort et al., 2012; Roelfsema et al., 2002). The present model went beyond these previous studies by using harder segmentation tasks and a neural network training scheme with a loss function for the segmentation outcome. The training procedure resulted in internal representations that resemble, to some extent, those of the primate brain, without the need to handcraft them. A related approach (Lei et al., 2021), carried out semantic segmentation of digits of the MNIST data set. Our work goes one step further by examining texture-defined figure-ground stimuli, and by implementing a biologically plausible form of top-down feedback. Indeed, in our model feedback was modulatory, because the activity was gated by feedforward activation and the model then compared the feedback activity to the feedforward activation, to determine the magnitude of the segmentation signal (Figure 1D). Our model was able to segregate figures from the background based on the difference in the orientation of texture elements and illustrated how top-down attention amplifies figure-ground signals, just as is observed in the visual cortex of monkeys (Poort et al., 2012). Furthermore, we evaluated the effect of feedback connections on the tuning of units, providing potential novel predictions, which can be tested in future neurophysiological experiments, e.g. the increase in shape selectivity and surround suppression due to feedback. A limitation of the present approach is that the network only used a feedforward and a feedback pass, but did not include full recurrence, even though the segmentation results are likely to further improve object-recognition, which in turn, may further refine the segmentation results, especially for shallower networks (Seijdel et al., 2020). Future research is needed to close this gap.

Our work focused on the relation between ANNs and mechanisms for image parsing in the brain. The aim therefore differed from that of ANN studies that focus on improving the state-of-the-art using methods without considering biological plausibility. Knowledge of brain-like mechanisms for vision may inspire improvements in the performance of ANNs in several ways. For example, the inclusion of feedback connections in ANNs does not only increase the similarity with the brain computations (Kietzmann et al., 2019; Kubilius et al., 2018), but improves classification accuracy and noise resistance (Jarvers and Neumann, 2019; Spoerer et al., 2020, 2017; Yan et al., 2019). Similarly, the inclusion of horizontal connections in the network, which are present between neurons in the visual cortex, aids in solving complex perceptual grouping tasks (Kim et al., 2019; Linsley et al., 2020, 2018). Hence, brain processes can inspire improvements in ANNs, levering an intense cross-fertilization between neuroscience and artificial intelligence (Doerig et al., 2022).

Overall, we provided a proof-of-concept demonstration that modulatory feedback are sufficient to produce accurate object segmentations of attended shapes. Our model represents a minimal architecture for object parsing in the brain, based on what we know about the roles of feedforward and feedback connections (Keller et al., 2020; Self et al., 2013; van Kerkoerle et al., 2017). We expect that future models will further improve our understanding of these mechanism, for example for more ecological stimuli to elucidate behavioral (Fowlkes et al., 2007; Neri, 2017, 2014; Zuiderbaan et al., 2017) and neural (Hesse and Tsao, 2016; Papale et al., 2018; Williford and von der Heydt, 2016) signatures of object segmentation in natural scenes. Future studies might also expand the biologically plausibility of neuronal networks for object-based attention by including horizontal connections and brain-like learning rules (Pozzi et al., 2020).

## Acknowledgments

We thank Jasper Poort for providing the code and pre-processed data from his experiment, and Alexander van Meegen for his early contribution to the model implementation. This work was supported by NWO (NWO-OCENW.KLEIN.178, Crossover Program 17619 “INTENSE” and “DBI2”, a Gravitation program of the Dutch Ministry of Science, Education and Culture; Open Competition Domain Science – XS, number OCENW.XS22.2.097), the European Union FP7 (ERC grant number 339490 ‘Cortic_al_gorithms’), Horizon Europe (ERC advanced grant 101052963 “NUMEROUS”; grant agreement 899287 “NeuraViper”), the Human Brain Project (agreement no. 945539, “Human Brain Project SGA3”) and *Agence Nationale de la Recherche* (ANR) within *Programme Investissements d’Avenir, Institut Hospitalo-Universitaire* FOReSIGHT (ANR-18-IAHU-0001).

## Notes

### Competing Interest Statement

The authors have declared no competing interest.

### Summary of Updates

-added supplementary information

