## Supplemental information for "Modulatory feedback determines attentional object segmentation in a model of the ventral stream"

### Supplementary methods

*RF position and the tuning of network units*

To examine the tuning of network units to space, orientation, border-ownership and shape and to assess surround suppression, we monitored their activity while varying the stimuli systematically, as would be done in electrophysiology experiments. We determined the RF of each unit using a light-spot mapping (e.g., Poort et al., 2016). We recorded responses from a set of ON and OFF stimuli, where a single pixel was either white (ON) or black (OFF) against a gray background. We reconstructed the spatial RF of the neuron and determined the RF center, as the mean position of the most influential pixels.

We determined the orientation tuning with full-contrast Gabor gratings with 8 different orientations (22.5 degrees apart), 3 spatial frequencies (2.5, 5 and 10 cycles per stimulus) and 8 phases (45 degrees apart). Following (Williford and von der Heydt, 2016), the orientation tuning index (OI), going from 0 (no tuning) to 1, was defined as following:

$$\mathrm{OI} = \frac{R_{pref} - R_{orth}}{R_{pref} + R_{orth}}$$

Where $R_{pref}$ was the response to the preferred orientation, spatial frequency and phase, and $R_{orth}$ was the response to the orthogonal orientation with the preferred spatial frequency and phase.

To measure surround suppression, we recorded responses from a set of Gabor patches with increasing sizes (2 pixel steps). We centered the patch on the RF and used the preferred combination of orientation, spatial frequency and phase. The surround suppression index (SSI) varies between 0 (no suppression) and 1 and was computed as follows:

$$\mathrm{SSI} = \frac{R_{pref} - R_{full}}{R_{pref} + R_{full}}$$

Where $R_{pref}$ was the response to a Gabor stimulus with the preferred size and $R_{full}$ was the response to a grating that filled the entire image.

To measure border ownership tuning we placed the edge of a square on the center of the RF of each neuron. The squares had 64 possible orientations (rotated in steps of 5.6 degrees), three sizes ($Preferred\ size, 2 \times Preferred\ size, 4 \times Preferred\ size$) and two possible locations (i.e., they appeared on the right or on the left). As in (Zhou et al., 2000), the border-ownership index (BOI), going from 0 (no selectivity) to 1, was defined as follows:

$$\mathrm{BOI} = \frac{R_{same} - R_{diff}}{R_{same} + R_{diff}}$$

Where $R_{same}$ is the response to the preferred stimulus, and $R_{diff}$ was the response to the stimulus having the same properties of $R_{same}$ (i.e., square size, orientation and contrast polarity) but on the opposite side.

To determine shape selectivity, we presented 2,000 stimuli with a particular shape. We averaged the activity per shape and quantified shape-selectivity as the sparsity of the distribution. Following (Zoccolan et al., 2007), the class-shape index (CSI), going from 0 (no selectivity) to 1, computed as following:

$$\mathrm{CSI} = [1 - \frac{\sum({R_i}/{12})^2}{({\sum R_i}^2/{12})}] \Big/ [1 - (\frac{1}{12})]$$

Where $R_i$ is the response to $i^{th}$ class and 12 is the number of classes.

Finally, we visualized each unit's tuning with an ANN feature visualization method (e.g. (Olah et al., 2017)). Contrast and luminosity in the images in the main figures have been slightly changed to enhance visibility, the original ones can be seen in the Colab of the project (see Methods).

**Supplementary results**

*Tuning of units in the feedforward and feedback paths of the network*

Given the similarity between the dynamics of activity in V1m and monkey V1 (Fig. 3), we investigated the tuning of units, following the similar procedures that are used when studying real neurons. For each of the units in the three lowest areas (V1m, V2m and V4m) of our model, we computed the degree of selectivity to many stimulus properties for which tuning in the primate visual system is common (Fig. S1A,B) and we also computed an image (image insets in Fig. S1C,D) that best drives the unit (Bashivan et al., 2019; Papale et al., 2021; Walker et al., 2019). Many neurons in the visual cortex are selective for the orientation of edges inside the RF (a schematic of typical responses is shown in Fig. Fig S1. Network units have tuning properties resembling those of neuronsA, left). This tuning relies to a large degree on the selectivity of feedforward connections to V1 (Chung and Ferster, 1998; Jin et al., 2011;

Lien and Scanziani, 2013). For the model units, we also quantified orientation tuning using the orientation index (OI, see Supplementary Methods) by centering stimuli with different orientations on the RF. The OI is the ratio between the difference response between the preferred and orthogonal orientation and the sum of these two responses. Most units in our model exhibited consistent tuning to the orientation of image elements (blue curves in Fig. S1C,D) similar to what is observed in the brain. For instance, V1m neurons in the feedforward pathway had a heterogeneous distribution of OIs with a median of 0.34, which is similar to the distribution and median OI of 0.39 of V1 neurons (Ringach et al., 2002).

Another common property of visual neurons is surround suppression (Fig. S1B, red), where image elements in the surround of the RF suppress the activity of a neuron. This property is the result of a suppressive influence of nearby neurons in the same cortical region and top-down feedback from neurons in higher regions (Angelucci et al., 2017; Bijanzadeh et al., 2018). Some of the units in our model showed surround suppression (Fig. S1C-E, red), which we quantified with the surround suppression index (SSI, see Methods). Surround suppression was present in both the feedforward and the feedback pathway, but it was relatively weak. Interestingly, while only 5% of V1m neurons presented some degree of surround suppression in the feedforward pathway, all of them showed surround suppression in the feedback pathway. In V2m and V4m, ~85% of the units had a non-zero SSI. Nevertheless, the surround suppression in the feedback pathway of the model (Fig. S2) was weaker than in V1 of animals (Bair et al., 2003; Jones et al., 2001). There are multiple possible reasons for this discrepancy. For example, the model did not include horizontal connections so that surround suppression was mediated by feedback only, whereas horizontal connections in the visual cortex of animals are thought to contribute to surround suppression (Adesnik et al., 2012).

Some visual neurons code for border-ownership (Qiu et al., 2007; Williford and von der Heydt, 2016), a property related to figure-ground perception and to the role of top-down feedback. In natural images, boundaries usually belong to a figure that occludes the background. Border-ownership tuning means that neurons are selective for the side of boundary that belongs to a figure. A schematic of the response of a typical border-ownership neuron is shown in the right panel of Fig. S1A. This example neuron responded more vigorously if the edge in the RF belonged to a figure that was positioned to the right of the edge than if it belonged to a figure on the left. In our model, there were no units with border-ownership selectivity. In Figure 4C-E, the tuning of the units reflects the orientation of the edge through the center of the RF, in accordance with their orientation tuning. However, these tuning profiles are symmetric, which indicates that the units were not selective for the figural side of the edge.

What causes the absence of border-ownership selective cells in our model? It is of interest that another modeling study (Dedieu et al., 2021) obtained units with border ownership tuning. They used a cloned Markov random field model in a similar setting as ours because the model had to parse MNIST digits from a background with distracting image elements. This model used several mechanisms, such as “cliques” and “query training”, which do not translate as easily into neurobiological mechanisms as the units and training curriculum used by us. Unlike our simulations, this previous study included stimuli in which objects could occlude each other. Border ownership signals are crucial for the correct assignment of contours formed by occluding, overlying shapes, which need to be ignored during recognition of the underlying, occluded objects.

We next investigated the tuning to shapes, a form of selectivity that is common for neurons in higher visual brain areas. Many of these neurons respond more strongly when a specific shape is present, irrespective of low-level features such as the position or size of the shape (Zoccolan et al., 2007). Many units in our model showed shape tuning (Fig. S1C-E, green), quantified as the shape-class selectivity index (CSI), measuring the sparseness of the distribution of response elicited by the 12 different shape classes. Shape tuning was stronger in the feedback path of higher model areas, with a CSI of 0.002 in V1m (same as in feedforward V1m), a CSI of 0.07 in V2m (0.001 in feedforward V2m) and a CSI of 0.7 in the feedback pathway of V4m (0.071 in feedforward V4m).

*The influence of feedback connections on the tuning of units at different network levels*

We next compared the tuning of units in the feedforward and feedback pathways in V1m-V4m. Figure S2 shows the distributions of orientation selectivity (blue), surround suppression (red) and object shape (green) in the feedforward and feedback pathways. We first examined orientation selectivity and observed no significant differences between the feedforward and feedback pathways in V1m, V2m and V4m (all ps $> 0.05$, Bonferroni corrected, Wilcoxon signed rank test). This finding suggests that orientation tuning of model units is largely determined by the selectivity of the feedforward connections. Surround suppression was stronger in the feedback units of V1m and V2m ($p < 0.01$, Bonferroni corrected, Wilcoxon signed rank test; Fig. S2A), suggesting a feedback influence on surround suppression in these lower network regions. In V4m, surround suppression was weak overall.

Finally, we examined to tuning to the different classes of shapes that the network had been trained on. Interestingly, tuning to the trained shapes was stronger in the feedback than in the feedforward pathways of V2m and V4m (both ps < 0.001; Fig. S2B,C), but this effect did not reach significance in V1m. Finally, we found no significant correlation between different tuning properties that we quantified, neither across feedforward units nor across feedback units (Pearson's correlation: all ps > 0.05, Bonferroni corrected, t-test), e.g. the selectivity of a unit for shape did not predict the degree of surround suppression.

All the tuning properties of model units can be explored in a Google Colab: https://colab.research.google.com/drive/13O0-4uYq3l1tZupVn-NLP8C-WvakajCG?usp=sharing.

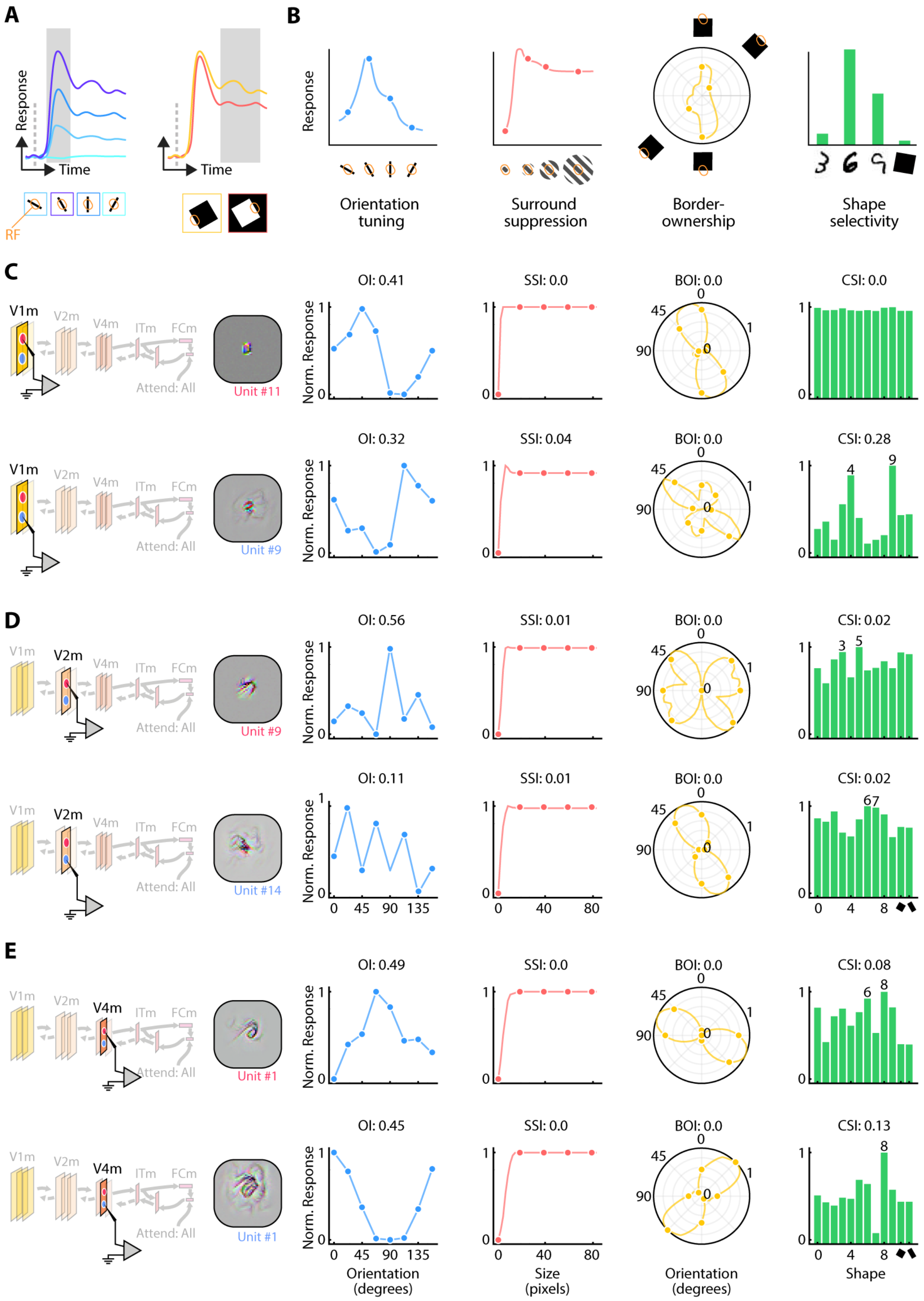


**Fig S1. Network units have tuning properties resembling those of neurons**

A) Schematic of responses of neurons in the visual cortex. Left: feedforward driven orientation tuning. The neurons respond more to some orientation than to others. Right: Border-ownership selectivity. The example neuron prefers stimuli for which the edge in the RF belongs to a figure on the right. B) Tuning of typical V1 neurons. From left to right: orientation tuning, surround suppression, i.e. neurons are suppressed if the size of the stimulus increases beyond the borders of the RF, border-ownership tuning and tuning to more complex shapes. C-E) Representative examples of the selectivity of feedforward (upper row) and feedback model units (lower row) in V1m (C), V2m (D) and V4m (E). The gray squares on the left show the most exciting image, which is a stimulus that best drives the units. Note the increasing tuning complexity from color-blob filters in V1m to object-part detectors in V4m (gray represents pixels that do not drive the cell, i.e. are outside of their RF). The other panels show the response to different orientations (blue; OI, orientation-index), stimulus sizes (red; SSI, surround suppression index), orientation and border-ownership tuning when one of the edges of a larger square is rotated within the RF (yellow; BOI, border-ownership index) and shape tuning (green; CSI, class-shape-selectivity index).

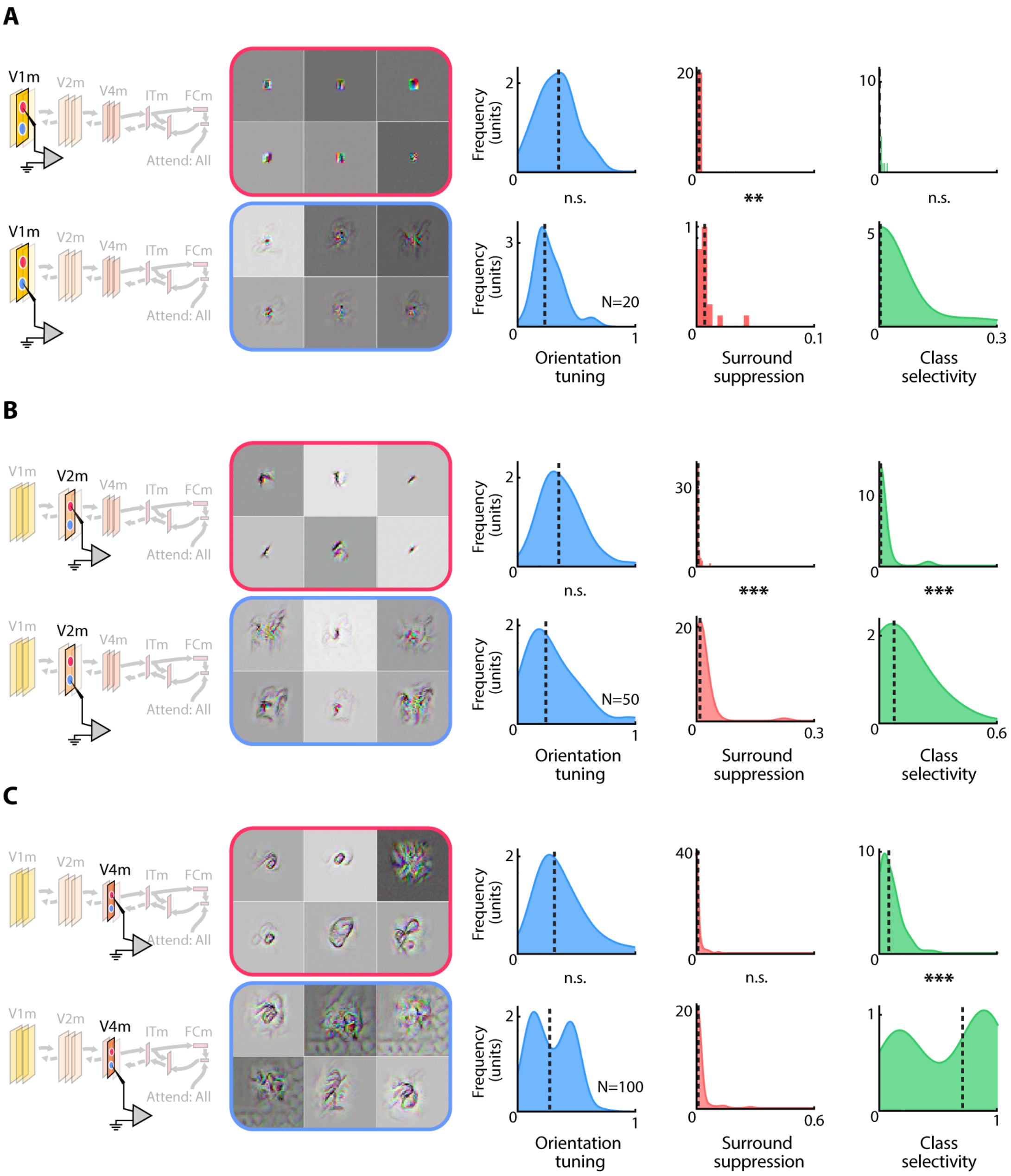


**Fig S2. Comparison of the tuning in the feedforward and feedback pathways**

Tuning of six example units (left) and distribution of orientation tuning (blue), surround suppression (red) and shape-selectivity (green) across all units in V1m (A), V2m (B) and V4m (C). In each panel, units of the feedforward are shown in the top row and units of the feedback pathway in the bottom row. The grey squares show the most exciting images, which maximally drive the units (gray indicates pixels outside the RF). We assessed significance of differences between the feedforward and feedback pathways with Wilcoxon signed rank tests (Bonferroni corrected). *n.s.*, $p > 0.05$; **: $p < 0.01$; ***: $p < 0.001$.
